# Exploring cell-specific miRNA regulation with single-cell miRNA-mRNA co-sequencing data

**DOI:** 10.1101/2020.10.14.340299

**Authors:** Junpeng Zhang, Lin Liu, Taosheng Xu, Wu Zhang, Chunwen Zhao, Sijing Li, Jiuyong Li, Nini Rao, Thuc Duy Le

## Abstract

**Background:** Existing computational methods for studying miRNA regulation are mostly based on bulk miRNA and mRNA expression data. However, bulk data only allows the analysis of miRNA regulation regarding a group of cells, rather than the miRNA regulation unique to individual cells. Recent advance in single-cell miRNA-mRNA co-sequencing technology has opened a way for investigating miRNA regulation at single-cell level. However, as currently single-cell miRNA-mRNA co-sequencing data is just emerging and only available at small-scale, there is a strong need of novel methods to exploit existing single-cell data for the study of cell-specific miRNA regulation.

**Results:** In this work, we propose a new method, *CSmiR* (<u>C</u>ell-<u>S</u>pecific <u>miR</u>NA regulation) to use single-cell miRNA-mRNA co-sequencing data to identify miRNA regulatory networks at the resolution of individual cells. We apply *CSmiR* to the miRNA-mRNA co-sequencing data in 19 K562 single-cells to identify cell-specific miRNA-mRNA regulatory networks for understanding miRNA regulation in each K562 single-cell. By analyzing the obtained cell-specific miRNA-mRNA regulatory networks, we observe that the miRNA regulation in each K562 single-cell is unique. Moreover, we conduct detailed analysis on the cell-specific miRNA regulation associated with the miR-17/92 family as a case study. Finally, through exploring cell-cell similarity matrix characterized by cell-specific miRNA regulation, *CSmiR* provides a novel strategy for clustering single-cells to help understand cell-cell crosstalk.

**Conclusions:** To the best of our knowledge, *CSmiR* is the first method to explore miRNA regulation at a single-cell resolution level, and we believe that it can be a useful method to enhance the understanding of cell-specific miRNA regulation.

## Background

As an abundant class of small, conserved and non-coding RNAs, microRNAs (miRNAs) play an important role in regulating gene expression through post-transcriptional repression or messenger RNA (mRNA) degradation [1]. In a cell, it is estimated that miRNAs can regulate the expression of up to one-third of the encoded human genes [2]. Such cellular effects of miRNAs influence a wide range of basic cellular functions, including cell proliferation, cell differentiation, and cell death [3]. Just as each individual cell is unique in the context of its microenvironment, miRNA regulation would tend to be unique in each individual cell accordingly. Previously, based on bulk RNA sequencing expression data from large populations of cells, many computational methods have been developed for exploring miRNA regulation [4, 5], but at the resolution of groups of cells. This may have obscured the heterogeneity of miRNA regulation across individual cells within these populations. Fortunately, single-cell RNA sequencing technology has now provided us the opportunity to study miRNA regulation at the single-cell level.

To investigate miRNA regulation at the single-cell level, Wang *et al*. [6] used a half-cell genomics approach to generate single-cell miRNA-mRNA co-sequencing expression data of 19 K562 half cells, and then applied Pearson correlation method to identify miRNA targets. By using the half-cell genomics method, a single cell is lysed and the lysate is split evenly into two half-cell fractions. Then, each half-cell fraction can be used for either miRNA or mRNA transcriptome sequencing. They have found that miRNA expression variability alone may cause non-genetic intercellular heterogeneity. However, the identification of the miRNA targets by their work was in the grouped 19 K562 half cells rather than individual K562 half cells, consequently ignoring the heterogeneity of miRNA regulation between single-cells. To investigate the heterogeneity of miRNA regulation between different single-cells, it is necessary to explore cell-specific miRNA regulation (i.e. one miRNA regulatory network for one cell).

Although single-cell miRNA-mRNA co-sequencing data is emerging, the number of single-cells included in each single-cell dataset is still small mainly due to the lack of mature single-cell RNA sequencing technology for genome-wide profiling of both mRNAs and miRNAs [7]. To explore cell-specific miRNA regulation using single-cell miRNA-mRNA co-sequencing data, in this work, we adapt the cell-specific network (*CSN*) method proposed in [8] to infer cell-specific miRNA-mRNA regulatory networks. Given a single-cell gene expression data set including *g* genes and *n* cells, *CSN* infers *n* cell-specific networks. Each cell-specific network is an undirected gene association network, and consists of *g* nodes corresponding to *g* genes and the edges representing undirected gene-gene associations. *CSN* uses a statistic (see Eq. (2) in the “Methods” section) to calculate the strength of a gene-gene association in each cell. To identify the gene-gene associations in each cell by using the statistic, *CSN* takes a one-sided hypothesis test. The null hypothesis is that two genes are independent in cell *k*, and the alternative hypothesis is that two genes are associated with each other in cell *k*. If the statistic of a gene-gene association in cell *k* is larger than a significant level (e.g. 0.01), the gene-gene association in cell *k* exists. Although *CSN* can infer cell-specific gene regulatory networks consisting of cell-specific gene-gene associations, it can’t be directly utilized to identify cell-specific miRNA regulatory network as described below.

To explore cell-specific miRNA regulation, our method *CSmiR* extends *CSN* from the following two aspects. Firstly, *CSN* is only applicable to single-cell gene expression data with more than 100 single-cells. To address the issue, we introduce pseudo-cells to enlarge the number of single-cells in a single-cell miRNA-mRNA co-sequencing data set with less than 100 single-cells. Secondly, *CSN* is developed to infer all types of gene-gene interactions from single-cell RNA sequencing data. For single-cell miRNA-mRNA co-sequencing data, we focus on identifying the interactions between miRNAs and mRNAs rather than all the types of interactions (including miRNA-miRNA, miRNA-mRNA and mRNA-mRNA interactions).

We have applied the proposed *CSmiR* method to single-cell miRNA-mRNA co-sequencing expression data across 19 K562 half cells, and the analysis results indicate that *CSmiR* can help with the investigation of miRNA regulation at the resolution of individual cells.

## Results and discussion

### The miRNA regulation in each K562 cell is unique

As discussed above, to investigate cell-specific miRNA regulation using single-cell miRNA-mRNA co-sequencing data with a small number of samples, we propose to interpolate pseudo-cells to the data before inferring the miRNA-mRNA interactions of interest (see the “Methods” section). Accordingly, as shown in Fig. 1, our proposed method *CSmiR* consists of three main components, Interpolating pseudo-cells by sampling with replacement, identifying cell-specific miRNA-mRNA regulatory networks, and downstream analysis with cell-specific networks. Following the workflow of *CSmiR*, we have identified 19 cell-specific miRNA-mRNA regulatory networks for the 19 K562 cells. In this section, we present the results on the investigation of the uniqueness of each K562 cell in terms of cell-specific miRNA-mRNA interactions and hub miRNAs.

**Figure 1.**
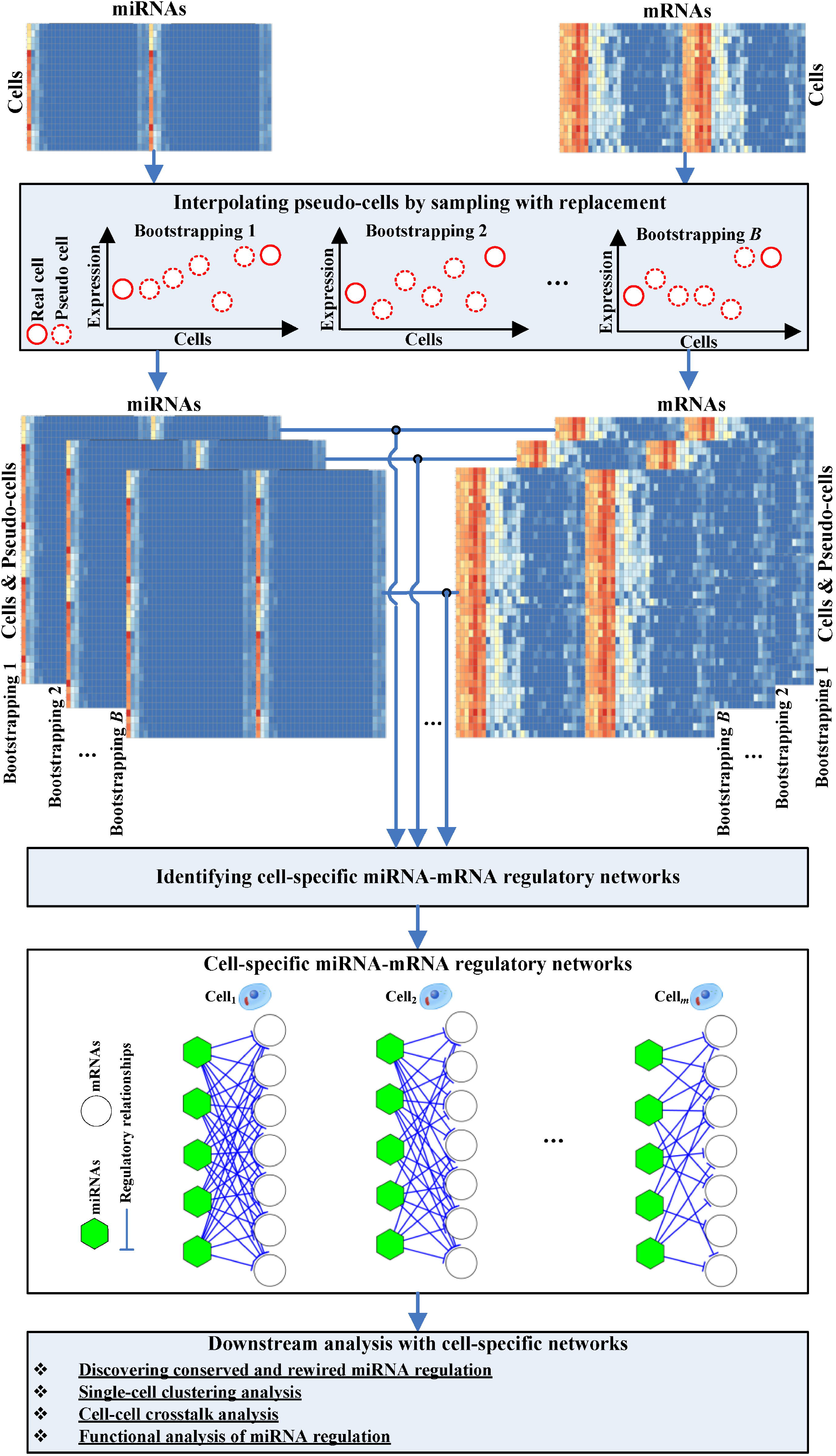
Workflow of *CSmiR*. For each pseudo-cell, we sample from the original dataset (i.e. the 19 single-cells uniformly with replacement) to generate it. Based on the *B* bootstrapping datasets (matched miRNA and mRNA expression data in the single-cells of the original dataset and interpolated pseudo-cells), we identify *m* cell-specific miRNA-mRNA regulatory networks for the real *m* cells (one miRNA-mRNA regulatory network for one cell). Finally, we conduct downstream analysis with the identified *m* cell-specific miRNA-mRNA regulatory networks.

Firstly, we have investigated the identified cell-specific miRNA-mRNA regulatory networks and hub miRNAs in four aspects: i) the number of predicted cell-specific miRNA-mRNA interactions, ii) the percentage of validated cell-specific miRNA-mRNA interactions, iii) the percentage of CML-related cell-specific miRNA-mRNA interactions, iv) the percentage of CML-related hub miRNAs. In the case of the four aspects, the miRNA regulation is different in each of the 19 K562 cells (see Fig. S1 in Additional file 1). Furthermore, we have discovered that the percentage of conserved and rewired miRNA-mRNA interactions is 20.47% (529998 out of 2588860) and 30.44% (787993 out of 2588860), respectively, indicating that the miRNA-mRNA interactions are more likely to be rewired across K562 cells. In terms of the similarity of the miRNA-mRNA interactions between these cell-specific regulatory networks, the range of cell similarity is [0.61, 0.86]. As shown in Fig. 2A, the cell similarity between any pair of the 19 K562 cells is less than 90%. In addition, the percentage of conserved and rewired hub miRNAs is 0% (0 out of 138) and 26.09% (36 out of 138) respectively, indicating the hub miRNAs tend to be rewired across K562 cells. In terms of hub miRNAs in the cell-specific regulatory networks, the range of cell similarity is [0.33, 0.62]. As shown in Fig. 2B, the cell similarity between any pair of the19 K562 cells is less than 70%.

**Figure 2.**
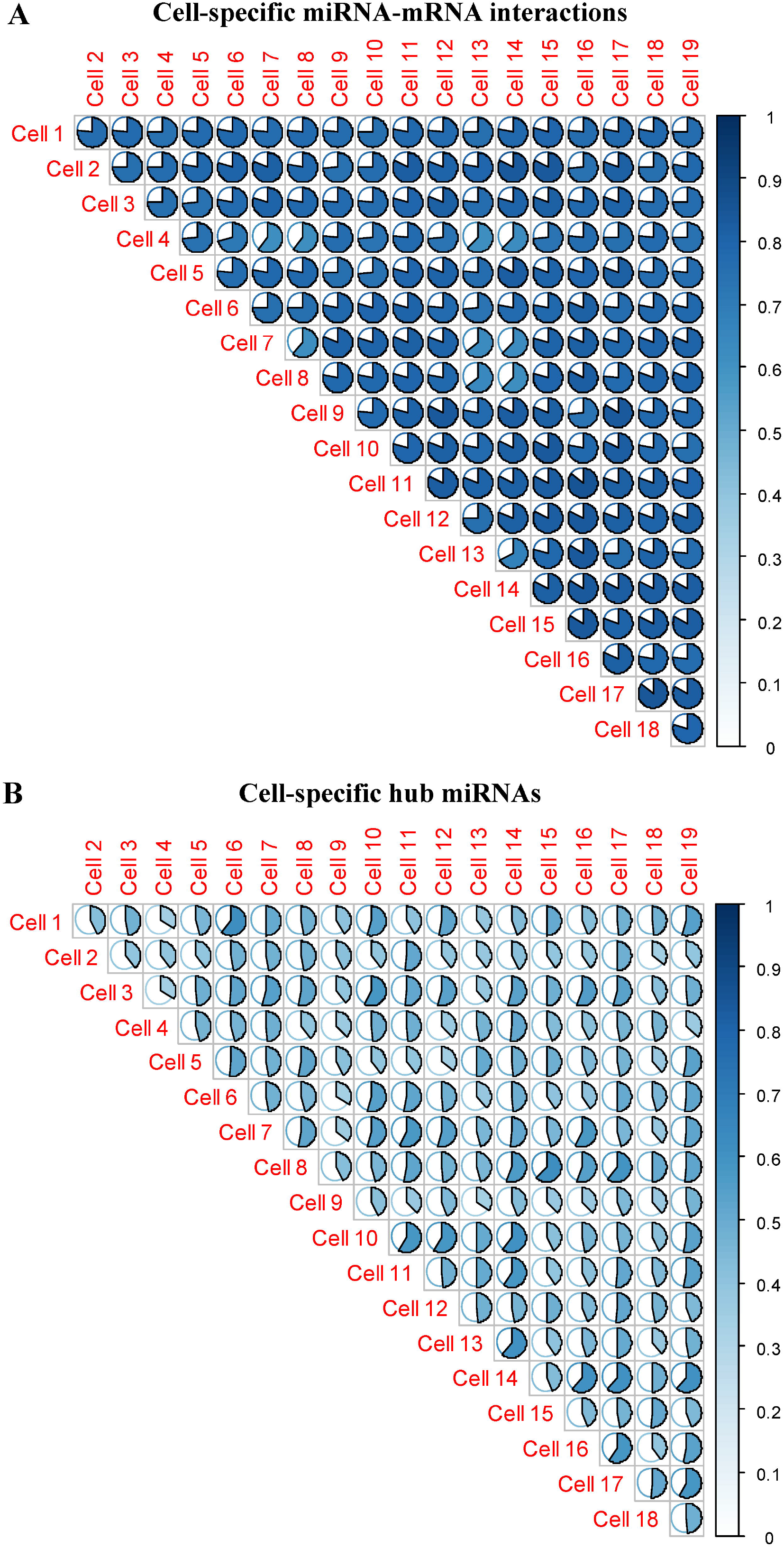
Single-cell similarity plot. (A) Similarity plot in terms of cell-specific miRNA-mRNA interactions. (B) Similarity plot in terms of cell-specific hub miRNAs. Colored areas indicate higher similarity between single-cells.

Comparing with the conserved miRNA regulation (conserved miRNA-mRNA interactions and hub miRNAs) across the 19 K562 single-cells, our work highlights a remarkable rewiring in the miRNA regulation (rewired miRNA-mRNA interactions and hub miRNAs) between the K562 cells, just like ‘on/off’ switches from cell to cell. The higher rewiring of miRNA regulation may be explained in part by the cell-specific expression of miRNAs and mRNAs, and it may be the reason of single-cell uniqueness. The detailed information of conserved and rewired miRNA-mRNA regulatory networks and hub miRNAs can be seen at https://github.com/zhangjunpeng411/CSmiR. Moreover, in terms of cell-specific miRNA-mRNA regulatory networks and cell-specific hub miRNAs, the above observations show that the miRNA regulation in any two different K562 cells are not completely the same, demonstrating the uniqueness of miRNA regulation in each cell.

### The miR-17/92 family regulation across K562 single-cells

To further understand the miRNA family regulation across K562 single-cells, we conduct a case study to investigate cell-specific regulation of the miR-17/92 family. The miR-17/92 family includes six members: miR-17 (miR-17-3p, miR-17-5p), miR-18a (miR-18a-3p, miR-18a-5p), miR-19a (miR-19a-3p, miR-19a-5p), miR-19b-1 (miR-19b-3p, miR-19b-1-5p), miR-20a (miR-20a-3p, miR-20a-5p) and miR-92a-1 (miR-92a-3p, miR-92a-1-5p). They play important roles in cell cycle, cell proliferation, cell apoptosis and other pivotal biological processes [9]. Previous studies [10–13] have also shown that the miR-17/92 cluster is in association with chronic myelogenous leukemia (CML). Out of the six members, miR-18a (miR-18a-3p and miR-18a-5p) with constant expression values across the 19 K562 single-cells is removed after data pre-processing. Hence in this section, we will focus on the regulation of the other five members (miR-17, miR-19a, miR-19b-1, miR-20a and miR-92a-1) from the miR-17/92 family.

To evaluate whether there is significant difference in the regulation of miR-17/92 family between each pair of the 19 K562 single-cells, we compare the distributions of the number of predicted targets, the distributions of the percentages of validated targets and the distributions of the percentages of CML-related targets of miR-17/92 family in different K562 single-cells using a two-sample Kolmogorov–Smirnov (KS) test [14]. The KS test is non-parametric, and can be used to assess whether the distribution of the number of predicted targets, the distribution of the percentages of validated targets or the distribution of the percentages of CML-related targets of miR-17/92 family in one K562 single-cell is significantly shifted compared with the distribution in another K562 single-cell. To estimate the distributions, we calculate the number of predicted targets, the percentage of validated targets and the percentage of CML-related targets of miR-17/92 family respectively in each K562 single-cell for each run of bootstrapping. As shown in Fig. 3A–3C, in the case of predicted targets, validated targets and CML-related targets, the regulations of miR-17/92 family between most of pairs of the 19 K562 single-cells are significantly different (*p*-value < 0.05). This result indicates that the regulation of miR-17/92 family is likely to be cell-specific. From Fig. 3D, the number of rewired targets of miR-17/92 family is larger than the number of conserved targets of them. This difference shows that the dominant miRNA regulation type (conserved or rewired) across cells may be rewired miRNA regulation. The detailed information of conserved and rewired targets associated with miR-17/92 family can be seen in Additional file 2.

**Figure 3.**
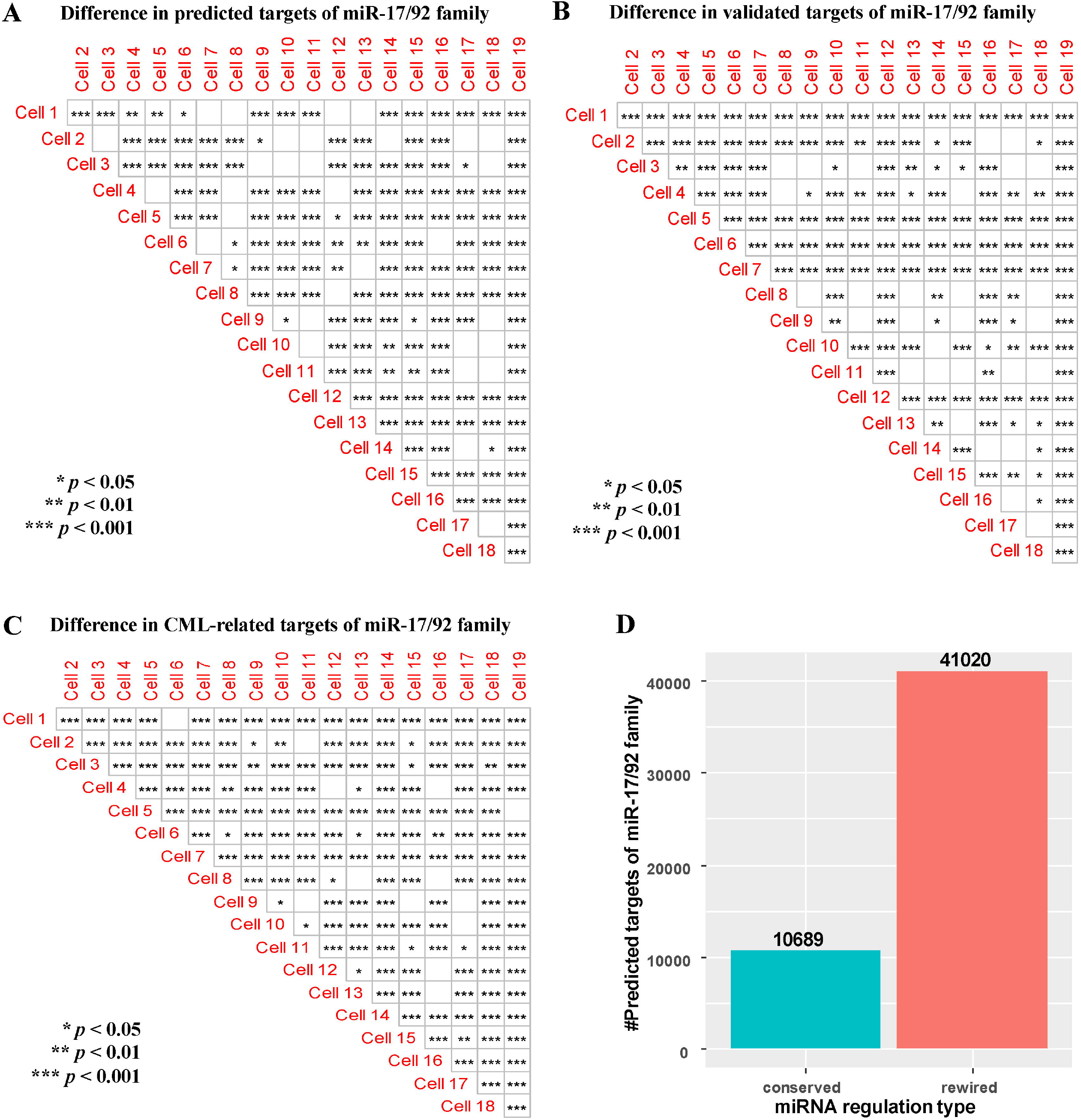
The miR-17/92 family regulation. (A) Difference in predicted targets of miR-17/92 family. (B) Difference in validated targets of miR-17/92 family. (C) Difference in CML-related targets of miR-17/92 family. (D) Number of conserved and rewired targets of miR-17/92 family. Empty square shapes denote *p*-values > 0.05.

Generally, through regulating target genes, miRNAs implement a specific biological function in the form of communities or modules. Therefore, based on the conserved and rewired miRNA-mRNA regulatory interactions associated with miR-17/92 family, we identify the conserved and rewired miRNA-mRNA modules associated with miR-17/92 family. We discover that most of the conserved and rewired miRNA-mRNA modules are significantly enriched in at least one term of Gene Ontology (GO), Kyoto Encyclopedia of Genes and Genomes Pathway (KEGG), Reactome, Hallmark or Cell marker (see Table S1 in Additional file 1). Several significant terms, e.g. the GO biological process “cellular response to leukemia inhibitory factor”, KEGG pathway “Chronic myeloid leukemia”, Reactome pathway “Regulation of mitotic cell cycle” [15], Hallmark “HALLMARK_TGF_BETA_SIGNALING” [16] and Cell marker “Peripheral blood, Leukemia, Cancer stem cell”, are closely associated with leukemia. This result shows that the identified conserved and rewired miRNA-mRNA modules associated with miR-17/92 family are functional modules. The detailed enrichment analysis results of conserved and rewired miRNA-mRNA modules can be seen in Additional file 3.

### *CSmiR* provides a novel strategy for clustering single-cells

Existing methods for clustering single-cells are mainly based on cluster analysis of single-cell RNA expression data. Different from these methods, we propose to cluster single-cells based on the interaction similarity and hub miRNA similarity as mentioned in the “Methods” section. We compare the proposed clustering method with the result of the clustering based on the Euclidean distance (normalized to the range of [0, 1]) between cells calculated using the expression data of single-cells (see the “Methods” section).

As shown in Fig. 4, we use hierarchical clustering to perform clustering analysis of the 19 K562 single-cells based on interaction similarity (Fig. 4A) and hub miRNA similarity (Fig. 4B) respectively, in comparison with the clustering based Euclidean distance (Fig. 4C). The clustering results differ due to different similarity/distance measures used. However, our method (either using the interaction similarity or hub miRNA similarity) gives rather distinct clusters, whereas the conventional cluster analysis directly based on the difference in gene expression does not produce any clear clusters. This result can be explained by previous studies [17, 18] showing that gene regulatory networks are more ‘stable’ than gene expressions to characterize the status of the biological process or cell. Although our clustering analysis results should be further validated by wet-lab experiments, *CSmiR* provides a novel strategy to help biologists discover clusters of cells which may indicate novel cell subtypes.

**Figure 4.**
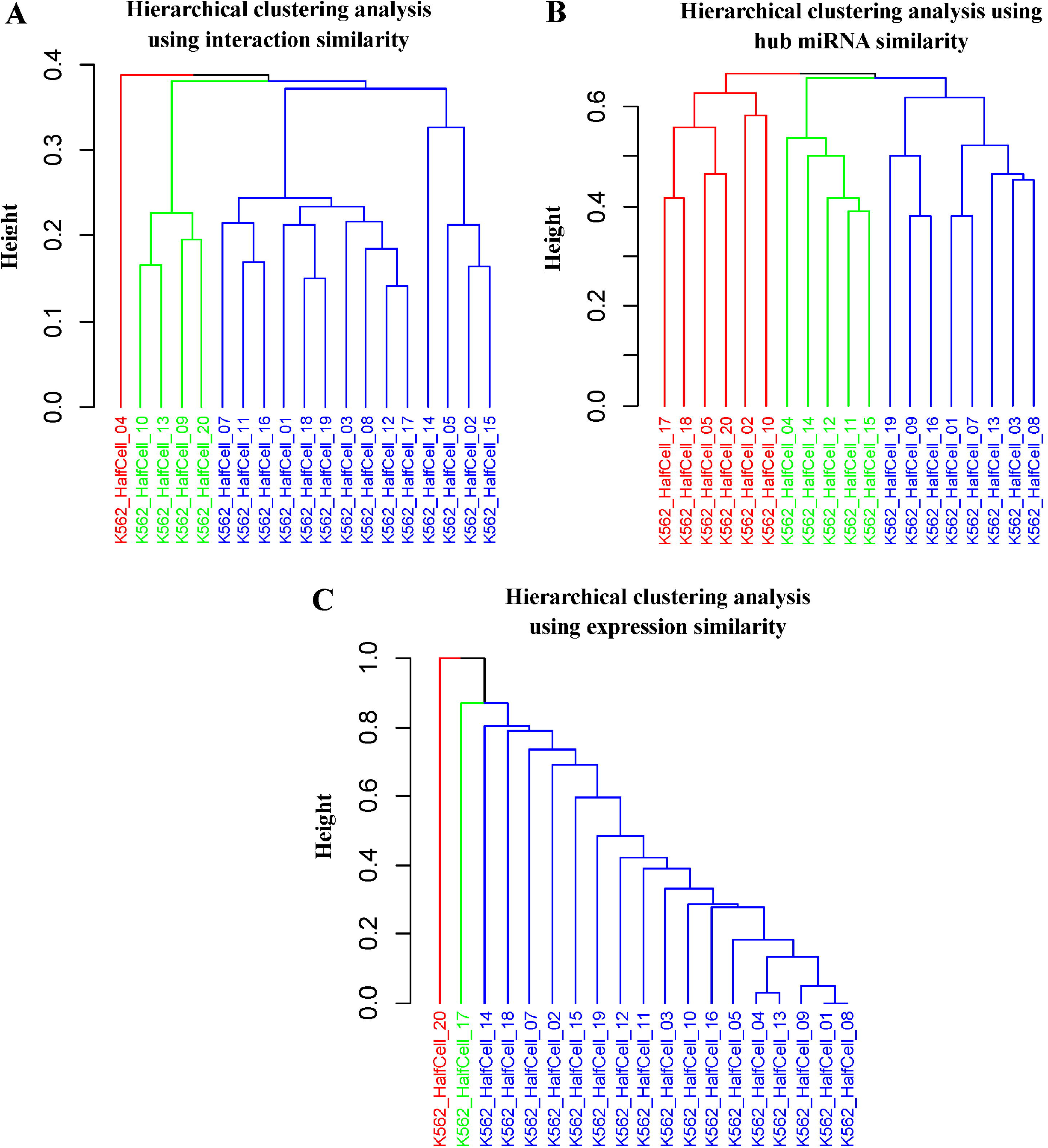
Hierarchical cluster analysis of the 19 K562 single-cells. (A) Hierarchical cluster analysis by using interaction similarity. (B) Hierarchical cluster analysis by using hub miRNA similarity. (C) Hierarchical cluster analysis by using expression similarity. Each color denotes a cluster.

### *CSmiR* helps to understand cell-cell crosstalk

It is known that cell-cell communication or crosstalk is crucial for multicellular organisms (i.e. human) because it allows multiple cells to communicate and coordinate to perform important life activity [19, 20]. Here, if the similarity value between cell*_i_* and cell*_j_* is larger than the median similarity value, cell*_i_* and cell*_j_* have a crosstalk relationship. In terms of interaction similarity or hub miRNA similarity, we assemble the cell-cell crosstalk relationships to generate cell-cell crosstalk network. Based on the interaction and hub miRNA similarity matrices, we obtain two cell-cell crosstalk networks (details in Additional file 4).

By analyzing the identified cell-cell crosstalk networks, we can understand which cells frequently communicate with other cells. We call these frequently communicated cells as hub cells or active cells. Similar to identifying cell-specific hub miRNAs, we also regard the top 20% of cells in terms of node degrees in each cell-cell crosstalk network as hub cells. These hub cells may act as pivots to link different subtypes of K562 single-cells (see Table S2 in Additional file 1). Moreover, we can also understand which cells tend to form a module in the process of communication. By using the Markov Clustering Algorithm (MCL) [21] implemented in the *miRspongeR* R package [22], we identify cell-cell crosstalk modules from the identified cell-cell crosstalk networks. For each module, the number of K562 single-cells is at least 3. We have discovered that most of the K562 single-cells only form a single module to communicate with each other (see Table S3 in Additional file 1). This observation can be explained that the 19 K562 single-cells used are phenotypically identical, and most of them are more likely form a module in cell-cell crosstalk.

## Conclusions

It is well known that miRNA regulation is essential to a wide range of important biological processes, including RNA silencing, transcriptional regulation of gene expression, cellular functions, signaling pathways and human cancers. Previous studies [23–25] have shown that miRNA regulation is condition-specific, implying that the miRNA regulation is cell-specific even these single-cells are phenotypically identical. Fortunately, single-cell RNA sequencing technology provides us an opportunity to gain insights into miRNA regulation at single-cell level. In this work, we have proposed *CSmiR*, a novel method to construct cell-specific miRNA-mRNA regulatory networks for each single-cell and use the networks to investigate cell-specific miRNA regulation. When identifying cell-specific miRNA-mRNA regulatory networks, since the cell-specific miRNA-mRNA regulatory networks are identified only from single-cell miRNA-mRNA co-sequencing expression data without using prior knowledge, *CSmiR* is an unsupervised method.

Our proposed method can be enhanced in several areas. Firstly, the identified cell-specific miRNA-mRNA networks are all correlation networks. Actually, to uncover miRNA causal regulation in single-cells, it is our future plan to identify cell-specific miRNA causal regulatory networks. Secondly, to improve the accuracy of the predicted cell-specific miRNA-mRNA regulatory networks, we can incorporate putative miRNA-mRNA binding information as prior knowledge into *CSmiR*. Finally, the miRNA regulation can be generally classified into two types: miRNA-directed regulation and miRNA-indirected regulation. In this work, we only consider the type of miRNA-directed regulation where miRNAs directly regulate the expression of mRNAs, and have not considered the type of miRNA-indirected regulation where miRNAs act as mediators to involve in gene regulation. According to the competing endogenous RNA (ceRNA) hypothesis [25], miRNAs act as mediators to involve in the crosstalk between different RNA transcripts (e.g. mRNAs, transcribed pseudogenes, circular RNAs and long noncoding RNAs). We also plan to infer cell-specific miRNA sponge interaction networks in future.

Although *CSmiR* can be improved as suggested above, it provides a new way to explore the heterogeneity of miRNA regulation in each single-cell. Especially, *CSmiR* can be applied in the study of germ cells or reproductive development [27], in which few cells could be profiled. We believe that *CSmiR* can be a useful method to speed up non-coding RNA (e.g. miRNA) research at single-cell level.

## Methods

In the following, we will describe the details about the single-cell miRNA-mRNA co-sequencing data used, interpolating pseudo-cells in small-scale single-cell transcriptomics data, the identification of cell-specific miRNA-mRNA regulatory networks, and subsequent analysis of the identified single-cell miRNA regulation.

### Single-cell miRNA-mRNA co-sequencing data

We obtain matched miRNA and mRNA co-sequencing expression data in 19 half K562 cells from Gene Expression Omnibus (GEO, https://www.ncbi.nlm.nih.gov/geo/) with accession number GSE114071. The K562 cells used are the first human chronic myelogenous leukemia (CML) cell line. For the duplicate miRNAs or mRNAs with the same gene symbols in the dataset, we compute the average expression values of them as their final expression values. Since gene expression variability may be a reason of non-genetic cell-to-cell heterogeneity [6], as a feature selection, we remove all the miRNAs and mRNAs with constant expression values (the standard deviation of their expression values in all single-cells is 0) across the 19 half K562 cells. The matched miRNA and mRNA expression data are then pre-processed by using log _2_ (*x* + 1) transformation. As a result, we have the matched expression profiles of 212 miRNAs and 15361 mRNAs in the 19 half K562 cells.

### Interpolating pseudo-cells in small-scale single-cell transcriptomics data

When the number of samples in a dataset is small, it is not guaranteed that a good representation of the population can be inferred from the data. It is required in [8] that when applying the *CSN* method, to estimate the association of each miRNA-mRNA pair, the number of cells in the single-cell transcriptomics dataset used should be more than 100. Since the proposed *CSmiR* method is adapted from the *CSN* method, for small-scale single-cell transcriptomics dataset like the one with 19 K562 half cells, it is necessary to enlarge the number of cells.

After interpolating pseudo-cells into original single-cell transcriptomics data, the main challenge is that the distribution of each gene (miRNA or mRNA) and joint distribution of each miRNA-mRNA pair will not be changed. To tackle this problem, we need to guarantee that the proportion of each cell type in the interpolated pseudo-cells is the same as that in the real single-cells. That is to say, the cell-type of the interpolated pseudo-cells should be the same as that in the real single-cells, and the number of the interpolated pseudo-cells of each cell-type also increases with the same probability. Based on this, for each pseudo-cell, we sample from the original single-cell transcriptomics data, i.e. the 19 single-cells uniformly with replacement, to generate it. To meet the requirement of having at least 100 single-cells, the number of interpolated pseudo-cells between two adjacent half K562 cells is set to 5. Here, two K562 single-cells with adjacent sample IDs (generated by half-cell genomics method) are regarded as adjacent single-cells. As a result, for each run of bootstrapping, we obtain the expression profiles of 212 miRNAs and 15361 mRNAs in 109 half K562 cells (including both real and pseudo half K562 cells). All the *B* bootstrapping datasets are used for subsequent analysis. In this work, the number of bootstrapping *B* is set to 100.

### Identifying cell-specific miRNA-mRNA regulatory networks

To reconstruct cell-specific miRNA-mRNA regulatory networks for real cells from the given single-cell dataset (including both real cells and interpolated pseudo-cells), it is necessary to construct a reliable statistic to evaluate the association between miRNAs and mRNAs. By using the statistic, the identified cell-specific miRNA-mRNA regulatory networks should be robust in the case of high dropout rate (also called technical noise from single-cell sequencing technology) and adding new cells (pseudo-cells in this work). Based on this, for each cell (including both real cells and interpolated pseudo-cells) in the given single-cell dataset, we apply the statistic used in the *CSN* method [8] to build a miRNA-mRNA regulatory network. In the case of high dropout rate and adding new cells, it is demonstrated that the *CSN* method is robust in identifying cell-specific networks. Therefore, when building the network, we adapt the *CSN* method for the discovery of miRNA-mRNA regulation. Specifically, for each miRNA-mRNA pair *miR_r_* and *mR_t_* in cell *k*, we evaluate the association between the miRNA and mRNA using the statistical test as described in the following.

To estimate the association between *miR_r_* and *mR_t_* in cell *k*, the *CSN* method draws a scatter diagram using the expression values of *miR_r_* and *mR_t_*. As shown in Fig. 5, *r_k_* and *t_k_* denote expression values of *miR_r_* and *mR_t_* in cell *k* respectively, and the medium, light and dark grey boxes represent the neighborhood of *r_k_*, *t_k_* and (*r_k_*, *t_k_*) respectively. The number of points in the medium, light and dark grey boxes are *n_r_*^(*k*)^, *n_t_*^(*k*)^ and *n_rt_*^(*k*)^ respectively. Then we construct the statistic, 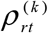 as:

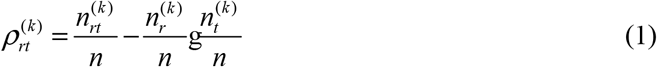

where *n* is the total number of cells in the given dataset, 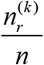 and 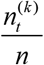 are the marginal probabilities of the expression levels of *miR_r_* and *mR_t_* respectively (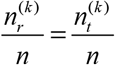 as empirically suggested by the *CSN* method), and 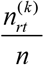 is the joint probability of *miR_r_* and *mR_t_*.

**Figure 5.**
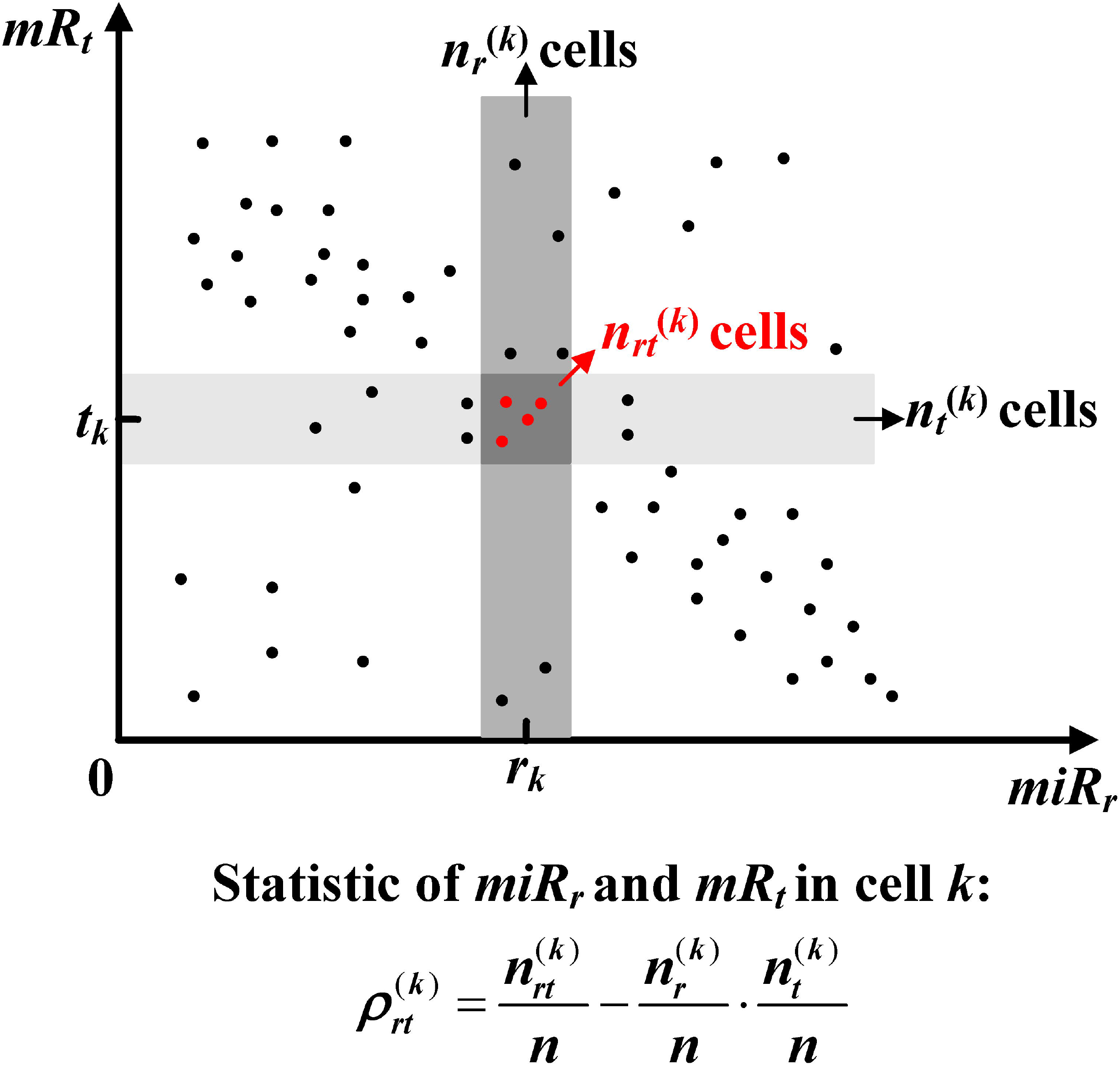
Statistic model for regulation between *miR_r_* and *mR_t_* in cell *k*. In the scatter diagram, *r_k_* and *t_k_* denote expression values of *miR_r_* and *mR_t_* in cell *k* respectively. The medium and light grey boxes denote the neighbourhood of *r_k_* and *t_k_*, respectively. The dark grey box (the intersection between the medium and light grey boxes) is the neighbourhood of (*r_k_*, *t_k_*). The number of points in the medium, light and dark grey boxes is *n_r_*^(*k*)^, *n_r_*^(*k*)^ and *n_rt_*^(*k*)^ respectively.

It has been proved in [8] that 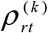 approximately follows a normal distribution, and the normalized statistic 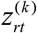 is:

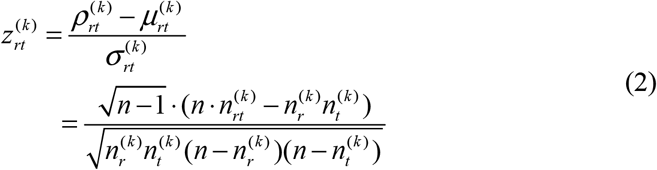

where 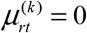 and 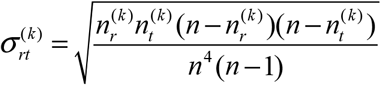 are the mean value and standard deviation of 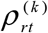, respectively. 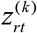 obeys standard normal distribution with mean value of 0 and standard deviation of 1. Each 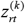 value corresponds to a *p*-value for evaluating the significance of the association between *miR_r_* and *mR_t_*. The smaller *p*-value indicates that the miRNA *miR_r_* and the mRNA *mR_t_* are more likely to be associated with each other in cell *k*. Here, the significant *p*-value cutoff is set to 0.01. For example, if we have a single-cell transcriptomics dataset containing 100 cells, the association between *miR_r_* and *mR_t_* in cell *k* is calculated as follows. Fig. 5 is the scatter diagram using the expression values of *miR_r_* and *mR_t_*. Then, we draw the two boxes near *r_k_* and *t_k_* based on the predetermined *n*_r_^(*k*)^ and *n_t_*^(*k*)^ 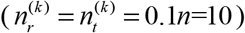. The value of 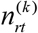 is 4 by counting the red points in the third box which is the intersection between the drawn two boxes. According to Eq. (1) and Eq. (2), the association 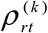 and normalized association 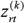 between *miR_r_* and *mR_t_* in cell *k* is 0.03 and 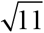. By using *pnorm* R function, the corresponding significance *p*-value of 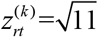 is 4.56E-04 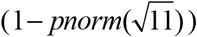 Given the significant *p*-value cutoff of 0.01, the miRNA *miR_r_* and the mRNA *mR_t_* are regarded as to be associated with each other in cell *k*.

Unstable estimation between the miRNA *miR_r_* and the mRNA *mR_t_* in cell *k* caused by small number of samples is a challenge to *CSmiR*. It is known that bootstrapping is a re-sampling technique used to obtain a reasonably accurate estimate of the population, and can be used to tackle the small sample problem. Therefore, to tackle this issue, we regard the median value of all the normalized associations 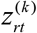 calculated in all the *B* runs of booststrapping as the final estimation of the asscociation between *miR_r_* and *mR_t_* in cell *k*. If the final association corresponds a small significance *p*-value (i.e. less than 0.01), the miRNA *miR_r_* and the mRNA *mR_t_* are associated with each other in cell *k*. As we are only interested in the miRNA-mRNA regulatory networks for each of the real cells (in the K562 dataset, they are the 19 real half K562 cells in the original dataset), at the end of this stage, we only keep the cell-specific miRNA-mRNA regulatory networks for the real cells. It is noted that each cell-specific miRNA-mRNA regulatory network is a bipartite graph where nodes are miRNAs and mRNAs and an edges is pointing from a miRNA to a mRNA.

### Downstream analysis with cell-specific networks

At the network level, it is known that gene regulatory network provides an insight into investigating gene regulation. In the same vein, the discovered cell-specific miRNA-mRNA regulatory networks in the previous step could also help to explore miRNA regulation. To explore cell-specific miRNA regulation, based on the identified cell-specific miRNA-mRNA regulatory networks, *CSmiR* conduct the following types of downstream analyses: i) Discovering conserved and rewired miRNA regulation, ii) Single-cell clustering analysis, iii) Cell-cell crosstalk analysis, and iv) Functional analysis of miRNA regulation.

#### Discovering conserved and rewired miRNA regulation

In a cell, the regulation of some miRNAs is “on” whereas the regulation of some miRNAs is “off” [27], indicated by having outgoing edges from the miRNAs or having no outgoing edges in the cell-specific network, respectively. It is possible that the regulation of some miRNAs is “on” in multiple cells and some miRNA regulations only maintain “on” in one cell. This “on/off state” phenomenon could help reveal the heterogeneity and commonality of miRNA regulation across different cells. Assuming that each cell is characterized by miRNA regulation, the conserved and rewired miRNA regulation across different cells can reflect the commonality and heterogeneity of cells, respectively. In this work, we discover conserved and rewired miRNA regulation in terms of both miRNA-mRNA regulatory network and hub miRNAs. Previous studies [28, 29] have shown that nearly 20% of the nodes in a biological network are regarded as essential nodes. The essential nodes in a biological network are subject to several topological properties (e.g. node degree) or biological relevance. Here, for simplicity, we select the top 20% of miRNAs based on node degrees in each cell-specific miRNA-mRNA regulatory network as hub miRNAs. Normally, if a miRNA-mRNA interaction or hub miRNA exists in more single-cells, the miRNA-mRNA interaction or hub miRNA tends to be more conservative. Here, the miRNA-mRNA interactions or hub miRNAs that are always “on” in at least 17 real K562 cells (~90%, generally ranked as a highly conservative level) are defined as conserved interactions or hubs, and the miRNA-mRNA interactions or hub miRNAs that are “on” in only one K562 cell are defined as rewired interactions or hubs. By assembling the conserved and rewired miRNA-mRNA interactions or hubs, we can obtain conserved and rewired miRNA-mRNA regulatory networks or hub miRNAs, respectively. These networks and hubs could provide insights into the heterogeneity and similarity of miRNA regulation across different single-cells.

#### Single-cell clustering analysis

Clustering single-cells based on single-cell RNA sequencing data is a fundamental task to understand tissue complexity, e.g. the number of subtypes [31]. In this paper, instead of directly using single-cell RNA sequencing data, we can use cell-cell similarity matrices for clustering single-cells, i.e. clustering cells based on their similarities on miRNA-mRNA interactions or hub miRNAs.

To reveal the heterogeneity of miRNA regulation across different cells, we investigate cell-cell similarity in terms of their miRNA-mRNA regulatory networks. The smaller the similarity between two single-cells is, the more heterogeneous they are.

We consider two types of similarities between two single-cells: the similarity on miRNA-mRNA interactions in their networks and the similarity on hub miRNAs in their networks. Following the similarity calculation method in [32], we calculate the interaction similarity and hub miRNA similarity using Eq. (3) below.

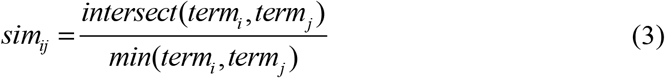

where *term_i_* and *term_j_* denote the numbers of interactions or numbers of hub miRNAs in the cell-specific miRNA-mRNA regulatory networks of cells *i* and *j*, respectively, *intersect*(*termi_i_*, *termj_j_*) denotes the number of miRNA-mRNA interactions or hub miRNAs common to the cell-specific miRNA-mRNA regulatory networks of cells *i* and *j*, and *min*(*termi_i_*, *termj_j_*) returns the smaller value out of *term_i_* and *term_j_*, i.e. the smaller value out of the numbers of miRNA-mRNA interactions or the numbers of hub miRNAs in the cell-specific miRNA-mRNA regulatory networks of cells *i* and *j*.

For comparison, we also calculate the similarity between two single-cells based on single-cell expression data. The normalized Euclidean distance *nor* _ *dis_ij_* between cells *i* and *j* is calculated as follows:

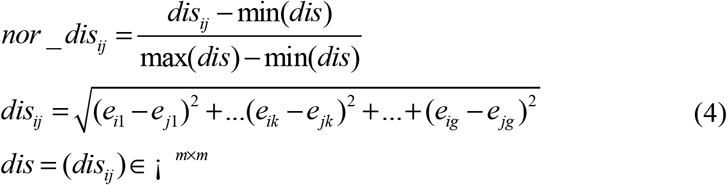

where *e_ik_* and *e_jk_* denote the expression levels of gene *k* in cells *i* and *j* respectively, *g* is the total number of genes (miRNAs and mRNAs), *m* is the number of real K562 single-cells.

After calculating the similarity and the distance between each pair of real half K562 cells, we obtain two similarity matrices *SI_m×m_* and *SH_m×m_* (where *m* is the number of real cells) in terms of cell-specific miRNA-mRNA interactions and cell-specific hub miRNAs respectively, and one distance matrix *dis_m×m_* in terms of single-cell expression data. Based on the similarity and distance matrices, we can conduct single-cell clustering analysis, e.g. hierarchical clustering analysis.

#### Cell-cell crosstalk analysis

In addition to single-cell clustering analysis, the similarity matrices can also be used for cell-cell crosstalk analysis. The cell-cell crosstalk is an indirect cell-cell communication and plays an important role in biological systems. For instance, cell-cell crosstalk can influence gene expression patterns [33], and involve in the development and regeneration of the respiratory system as well [34]. For each cell-cell pair, a higher similarity means sharing more number of miRNA-mRNA interactions or hub miRNAs between two cells. Previous studies [35, 36] have shown that miRNAs and their targets play important roles in cell signaling pathways. Therefore, when the shared miRNA-mRNA interactions or hub miRNAs involve in cell signaling pathways, a higher similarity between the cell pair implies that the two cells share more common cell signaling pathways and have a higher probability of signaling with each other (crosstalk). Based on this assumption, empirically, we use the median similarity value of all cell-cell pairs in the interaction or hub miRNA similarity matrix as the cutoff to evaluate whether two cells have crosstalk relationship or not. That is, if the similarity value between cell*_i_* and cell*_j_* is larger than the median similarity value, cell*_i_* and cell*_j_* have a crosstalk relationship. Following the empirical principle, we can evaluate whether each cell-cell pair has a crosstalk relationship or not. After assembling the cell-cell crosstalk relationships in terms of miRNA-mRNA interactions or hub miRNAs, we can obtain a cell-cell crosstalk network.

#### Functional analysis of miRNA regulation

To validate and apply the identified cell-specific miRNA regulatory networks, we also conduct functional analysis of miRNA regulation at both network and module levels. At the network level, we conduct functional validation of the cell-specific miRNA-mRNA regulatory networks by using third-party databases. Since there are no experimentally validated databases at single-cell level, we use two well-known experimentally validated databases named miRTarBase v8.0 [37] and TarBase v8.0 [38] at bulk-cell level for validation. Meanwhile, since the K562 cells used are closely associated with chronic myelogenous leukemia (CML), we collect a list of miRNAs and mRNAs associated with CML to investigate CML-related miRNA regulation. The list of CML-related miRNAs is from Human MicroRNA Disease Database HMDD v3.0 [39], and the list of CML-related mRNAs is from DisGeNET v5.0 [40], which is one of the largest publicly available collections of genes and variants associated to human diseases. We focus on identifying CML-related miRNA-mRNA pairs where the miRNAs and mRNAs individually are in the list of CML-related miRNAs and mRNAs.

At the module level, we discover miRNA-mRNA regulatory modules by using the *biclique* R package [41]. We consider each miRNA-mRNA regulatory module is a complete bipartite graph or a biclique, and the numbers of miRNAs and mRNAs in each module are at least 2 and 3, respectively. Here, a complete bipartite graph or a biclique is a special type of bipartite graph where every miRNA is connected to every mRNA. To understand potential biological implications associated with the identified miRNA-mRNA regulatory modules, we perform Gene Ontology (GO) [42], Kyoto Encyclopedia of Genes and Genomes (KEGG) [43], Reactome [44], Cancer hallmark [45], and Cell marker [46] enrichment analysis by using the *clusterProfiler* R package [47]. A GO, KEGG, Reactome, Hallmark or Cell marker term with adjusted *p*-value < (adjusted by Benjamini-Hochberg method) is regarded as a significant term. We also conduct CML enrichment analysis by using a hyper-geometric test to evaluate whether the miRNAs and mRNAs in each module are significantly enriched in CML or not. The significance *p*-value of each module enriched in CML is calculated as:

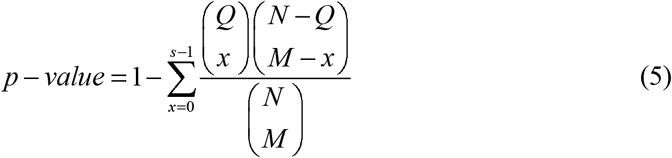

where *N* is the total number of genes (miRNAs and mRNAs) expressed in the dataset, *Q* represents the number of CML-related genes in the dataset, *M* is the total number of genes in each module, and *s* is the number of CML-related genes in each module. The cutoff of *p*-value is set as 0.05.

## Abbreviations

miRNA: microRNA
mRNA: messenger RNA
CSN: Cell-Specific Network
CSmiR: Cell-Specific miRNA regulation
GGI: Gene-Gene Interaction
MCL: Markov Clustering Algorithm
ceRNA: Competing Endogenous RNA
GEO: Gene Expression Omnibus
CML: Chronic Myelogenous Leukemia
GO: Gene Ontology
KEGG: Kyoto Encyclopedia of Genes and Genomes Pathway
KS: Kolmogorov-Smirnov

## Declarations

### Ethics approval and consent to participate

Not applicable.

### Consent for publication

Not applicable.

### Availability of data and materials

*CSmiR* is released under the GPL-3.0 License, and is available at https://github.com/zhangjunpeng411/CSmiR. Gene expression profiles used for *CSmiR* were accessed at Gene Expression Omnibus (https://www.ncbi.nlm.nih.gov/geo/). The lists of all the data used in this study are available in the additional files.

### Authors’ contributions

JZ, NR and TDL conceived the idea of this work. LL, TX and JL refined the idea. JZ designed and performed the experiments. TX, WZ, CZ and SL participated in the design of the study and performed the statistical analysis. JZ, LL, NR, TDL and JL drafted the manuscript. All authors revised the manuscript. All authors read and approved the final manuscript.

## Additional files

**Additional file 1 – Supplementary figure S1 and tables S1-S2.**

**Additional file 2 – Conserved and rewired targets associated with miR-17/92 family.**

**Additional file 3 – Enrichment analysis of conserved and rewired miRNA-mRNA modules associated with miR-17/92 family.**

**Additional file 4 – Cell-cell crosstalk networks.**

