## Additional file 1 for "Exploring cell-specific miRNA regulation with single-cell miRNA-mRNA co-sequencing data"

^1^Center for Informational Biology, School of Life Science and Technology, University of Electronic Science and Technology of China, China, ^2^School of Engineering, Dali University, China, ^3^UniSA STEM, University of South Australia, Australia, ^4^Institute of Intelligent Machines, Hefei Institutes of Physical Science, Chinese Academy of Sciences, China, ^5^School of Agriculture and Biological Sciences, Dali University, China

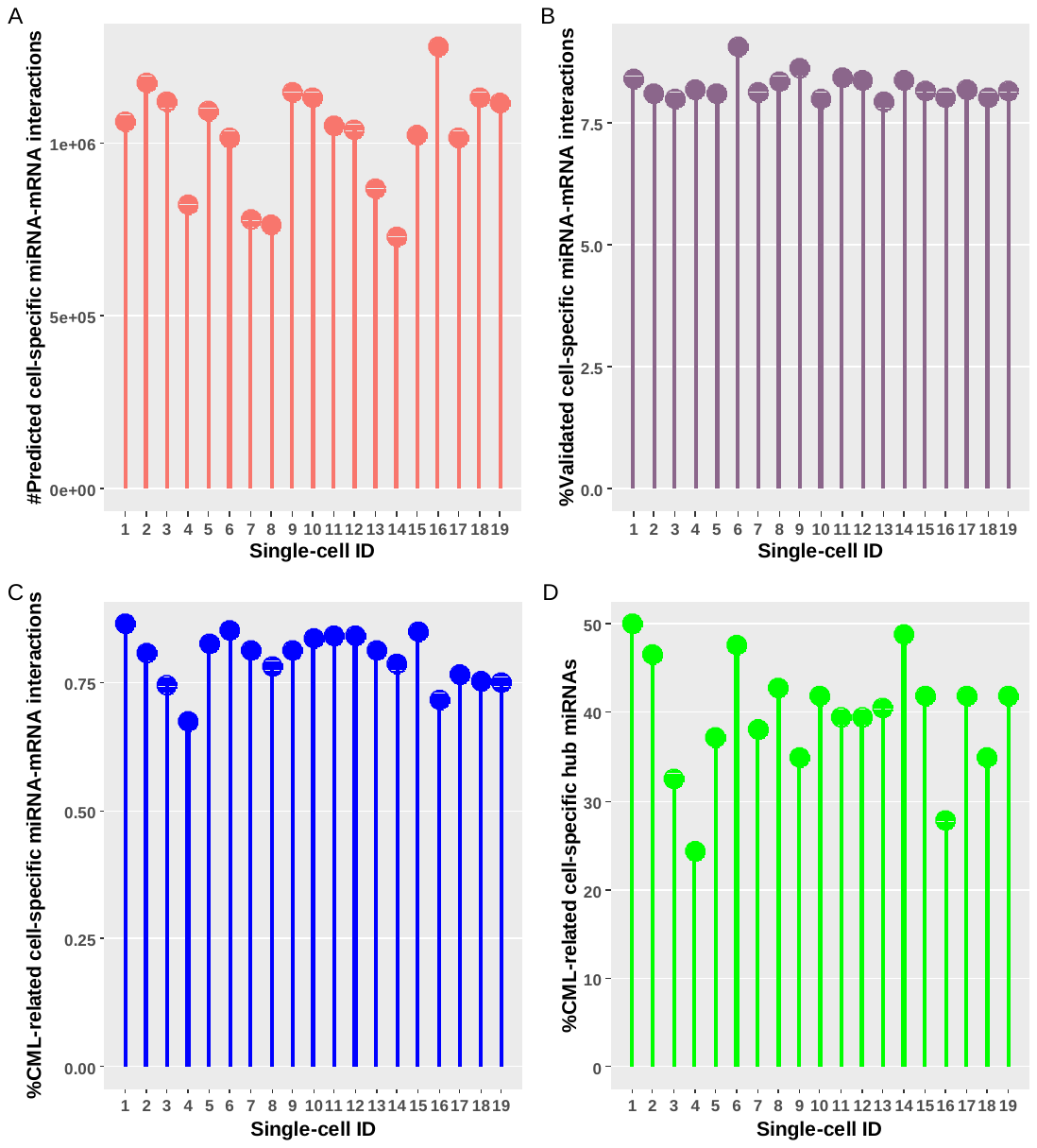

**Fig. S1** Cell-specific miRNA-mRNA interactions and hub miRNAs. (A) Number of predicted cell-specific miRNA-mRNA interactions. (B) Percentage of validated cell-specific miRNA-mRNA interactions. (C) Percentage of CML-related cell-specific miRNA-mRNA interactions. (D) Percentage of CML-related hub miRNAs.

**Table S1.** Enrichment analysis of conserved and rewired miRNA-mRNA modules associated with miR-17/92 family.

| Module type | ID | #GO | #KEGG | #Reactome | #Hallmark | #Cell marker | Enriched in CML or not? |
| --- | --- | --- | --- | --- | --- | --- | --- |
| Conserved | **1** | **0** | **0** | **0** | **0** | **0** | **No** |
|  | **2** | **83** | **14** | **13** | **2** | **0** | **No** |
|  | **3** | **1** | **0** | **0** | **0** | **15** | **No** |
|  | **4** | **0** | **0** | **0** | **0** | **0** | **No** |
| Rewired | **1** | **990** | **55** | **413** | **11** | **0** | **No** |
|  | **2** | **0** | **0** | **0** | **0** | **10** | **No** |
|  | **3** | **1** | **0** | **0** | **2** | **22** | **No** |
|  | **4** | **0** | **0** | **0** | **0** | **27** | **No** |
|  | **5** | **1618** | **110** | **630** | **16** | **0** | **No** |
|  | **6** | **1621** | **110** | **630** | **16** | **0** | **No** |
|  | **7** | **1626** | **112** | **626** | **16** | **0** | **No** |
|  | **8** | **1646** | **117** | **643** | **19** | **0** | **No** |
|  | **9** | **1029** | **62** | **428** | **11** | **0** | **No** |
|  | **10** | **1031** | **63** | **428** | **11** | **0** | **No** |
|  | **11** | **1045** | **69** | **441** | **12** | **0** | **No** |
|  | **12** | **1032** | **62** | **427** | **11** | **0** | **No** |
|  | **13** | **1062** | **69** | **440** | **12** | **0** | **No** |
|  | **14** | **1063** | **69** | **440** | **12** | **0** | **No** |
|  | **15** | **0** | **0** | **2** | **2** | **0** | **No** |
|  | **16** | **0** | **0** | **0** | **0** | **1** | **No** |

**Table S2.** Hub cells in each cell-cell crosstalk network. The cell-cell crosstalk networks are generated in terms of network similarity and hub miRNA similarity.

| Terms | Hub cells | Degree |
| --- | --- | --- |
| Network similarity | K562_HalfCell_12 | 17 |
|  | K562_HalfCell_16 | 15 |
|  | K562_HalfCell_18 | 14 |
|  | K562_HalfCell_13 | 13 |
| Hub miRNA similarity | K562_HalfCell_15 | 13 |
|  | K562_HalfCell_03 | 12 |
|  | K562_HalfCell_12 | 12 |
|  | K562_HalfCell_20 | 12 |

**Table S3.** The identified cell-cell crosstalk modules. The cell-cell crosstalk networks are generated in terms of network similarity and hub miRNA similarity.

| Terms | Module ID | K562 HalfCell ID |
| --- | --- | --- |
| Network similarity | 1 | 01, 02, 03, 05, 07, 08, 09, 10, 11, 12, 13, 14, 15, 16, 17, 18, 19, 20 |
| Hub miRNA similarity | 1 | 01, 02, 03, 04, 05, 07, 08, 09, 11, 12, 13, 14, 15, 16, 17, 18, 19, 20 |
